# SIRPα on mouse B1 cells restricts lymphoid tissue migration and natural antibody production

**DOI:** 10.1101/2020.05.13.092494

**Authors:** Katka Franke, Saravanan Y. Pillai, Mark Hoogenboezem, Marion J.J. Gijbels, Hanke Matlung, Judy Geissler, Hugo Olsman, Chantal Pottgens, Patrick J. van Gorp, Maria Ozsvar-Kozma, Yasuyuki Saito, Takashi Matozaki, Taco W. Kuijpers, Rudi W. Hendriks, Georg Kraal, Christoph J. Binder, Menno P.J. de Winther, Timo K. van den Berg

## Abstract

The inhibitory immunoreceptor SIRPα is expressed on myeloid and neuronal cells and interacts with the broadly expressed CD47. CD47-SIRPα interactions form an innate immune checkpoint and its targeting has shown promising results in cancer patients. Here, we report expression of SIRPα on B1 lymphocytes, a non-conventional subpopulation of murine B cells responsible for the production of natural antibodies. Mice defective in SIRPα signaling (SIRPα^ΔCYT^ mice) displayed an enhanced CD11b/CD18 integrin-dependent B1 cell migration from the peritoneal cavity to the spleen, local B1 cell accumulation, and enhanced circulating natural antibody levels, which was further amplified upon immunization with T-independent type 2 antigen. As natural antibodies are atheroprotective we investigated the involvement of SIRPα signaling in atherosclerosis development. Bone marrow (SIRPα^ΔCYT^>LDLR^−/−^) chimaeric mice developed reduced atherosclerosis accompanied by increased natural antibody production. Collectively, our data identify SIRPα as a unique B1 cell inhibitory receptor acting to control B1 cell migration, and imply SIRPα as a potential therapeutic target in atherosclerosis.

## Introduction

Signal regulatory protein alpha (SIRPα) is an inhibitory immunoreceptor known to be expressed on myeloid and neuronal cells. SIRPα interacts with the broadly expressed cell surface ligand CD47 present on most cells in the body, including both hematopoietic and non-hematopoietic cells (Barclay and van den Berg, 2014). Binding of CD47 to SIRPα generates intracellular inhibitory signals via immunoreceptor tyrosine-based inhibitory motifs (ITIM) in the cytoplasmic tail of SIRPα. Upon phosphorylation the SIRPα ITIM act to recruit and activate the tyrosine phosphatases SHP-1 and/or SHP-2, which inhibit tyrosine-phosphorylation-dependent signaling events and the resultant downstream cellular effector functions, including e.g. phagocytosis (Barclay and van den Berg, 2014). As such, the CD47-SIRPα axis forms an important innate immune checkpoint, with CD47 acting as so-called “don’t-eat-me” signal, which prevents the engulfment of healthy cells by myeloid cells (Matlung et al., 2017). However, aberrant cells, such as cancer cells, may also exploit this pathway by (over)expressing CD47 and thus escaping immune-mediated destruction. It is therefore not surprising that therapeutic targeting of the CD47-SIRPα checkpoint has been most intensively explored in the context of cancer. In fact, recent first in-human studies of agents interfering with this pathway demonstrate a favorable safety profile and promising therapeutic potential (Advani et al., 2018).

Based on their functions, anatomical location and phenotypical properties B lymphocytes can be divided into conventional B cells, also known as B2 cells, representing the majority of B cells, and into a smaller population of unconventional B1 cells. In mice, B1 cells are produced in the fetal liver before birth and afterwards reside mainly in the pleural and peritoneal cavities where they are maintained by self-renewal (Baumgarth, 2011). In addition, small proportions (<1%) of these cells can be found in spleen and bone marrow (Baumgarth, 2011; Choi et al., 2012; Smith and Baumgarth, 2019). B1 cells residing in body cavities have a limited capacity to produce natural antibodies. However, after stimulation, by e.g. LPS, they migrate to the secondary lymphoid tissues, such as the spleen, where they differentiate into plasma cells forming the major systemic source of natural antibodies. This conditional migration is governed by the CD11b/CD18 integrin (Waffarn et al., 2015). B1 cells that have arrived to the spleen gradually lose expression of CD11b/CD18 integrin, with hardly detectable levels after 6 days (Ghosn et al., 2008). Peritoneal B1 cells represent about 35-70% of all CD19^+^ cells present in the peritoneal cavity and can be further divided into B1a (CD19^+^CD11b^+^CD5^+^) and B1b (CD19^+^CD11b^+^CD5^−^) cells (Baumgarth, 2011). Unlike B2 cells, B1 cells in the spleen constitutively secrete natural antibodies, which are more promiscuous IgM antibodies targeting e.g. phospholipid and polysaccharide antigens, such as phosphorylcholine, phosphatidylcholine and lipopolysaccharide (Baumgarth, 2011). Notably, a large part of the natural IgM antibodies is directed against epitopes created through lipid peroxidation (so called oxidation-specific epitopes, OSE), expressed amongst others on apoptotic cells and modified lipoproteins (Miller and Tsimikas, 2013). Protective effects of natural antibodies against oxidized lipids have been well established in atherosclerosis (Binder et al., 2003; Caligiuri et al., 2007; Faria-Neto et al., 2006; Grasset et al., 2015), a chronic inflammatory disease characterized by accumulation of modified (oxidized) lipids in big and medium sized arteries (Libby et al., 2011). The atheroprotective capacity of IgM antibodies is explained by their binding to oxLDL, thereby preventing oxLDL uptake by macrophages, which as a consequence reduces foam cell formation and lesion development (Binder et al., 2003; Horkko et al., 1999). Additionally, natural antibodies are produced to promote clearance of apoptotic cells, which carry the same OSE as oxLDL (Grasset et al., 2015).

It is known that B1 cell responses are restricted by different inhibitory immunoreceptors expressed on these cells, including e.g. CD5 (Bikah et al., 1996), CD22 (Nitschke et al., 1997), Fc gamma receptor IIb (FcγRIIb) (Amezcua Vesely et al., 2012; Bagchi-Chakraborty et al., 2019), and Siglec-G (Ding et al., 2007; Hoffmann et al., 2007). CD5 has been strongly linked to inhibition of BCR signaling, which prevents unwanted self-reactivity of B1 cells (Hayakawa et al., 1983). B1 cells from mice lacking Siglec-G show a dramatic increase in Ca^2+^ flux upon anti-IgM treatment and increased natural antibody production, also suggesting role of Siglec-G in BCR signaling. All these receptors commonly exhibit their inhibitory functions through intracellular immunoreceptor tyrosine-based inhibitory motifs (ITIM), which upon tyrosine phosphorylation recruit and activate the cytosolic tyrosine phosphatases SHP-1 and/or SHP-2. In the case of FcγRIIb, the inositol phosphatases SHIP-1 and/or SHIP-2 play a prominent role as mediators of inhibitory signaling (Smith and Clatworthy, 2010).

Here we describe another inhibitory receptor, SIRPα, which is selectively expressed on B1 cells among B cell populations of mice. We demonstrate that, in contrast to other currently known inhibitory receptors, SIRPα on B1 cells negatively regulates their migration, B1 cell numbers in the spleen, and systemic natural antibody production, without directly affecting B1 cell activation. Mice lacking the cytoplasmic tail of SIRPα (SIRPα^ΔCYT^ mice) in their hematopoietic compartment are protected against atherosclerosis with increased natural antibody levels against oxidized lipids. This identifies SIRPα as a novel immunoinhibitory receptor on B1 cells with unique regulatory functions and potential for therapeutic targeting in atherosclerosis.

## Results

### SIRPα is expressed on B1 cells

The inhibitory immunoreceptor SIRPα is considered to be present selectively on neuronal cells as well as on myeloid cells in the hematopoietic compartment (Adams et al., 1998; Barclay and van den Berg, 2014). It is thought to be lacking from lymphoid cells, at least under steady state conditions (Sato-Hashimoto et al., 2011). However, a more detailed evaluation of B cell subsets revealed SIRPα expression on all B1 cells in the peritoneal cavity (PC) and on a subpopulation of B cells in the spleen (SP) of mice (Fig. 1A-C, Suppl. Fig. 1A, B). In particular, by using specific markers identifying B1a cells (i.e. CD19^+^CD5^+^CD11b^+^) and B1b cells (i.e. CD19^+^CD5^−^CD11b^+^) we could clearly demonstrate surface SIRPα expression on both of these PC B1 lymphocyte populations. Conventional PC B2 cells show much lower if any SIRPα expression. In the spleen we could detect expression of SIRPα only on a subset of B220^+^/CD19^+^CD43^+^CD23^−^ B1 cells. Relatively low levels of SIRPα staining were found on marginal zone B cells (B220^+^/CD19^+^CD43^−^CD23^−^) and minimal detectable expression was found on B220^+^/CD19^+^CD23^+^CD43^−^ follicular B cells. Consistent with other studies (Sato-Hashimoto et al., 2011) we could not observe any SIRPα expression on circulating B cells in mice and no expression on B2 cells from the bone marrow (Suppl. Fig. 1C). However, we could detect SIRPα on fetal liver B220^+^/CD19^+^CD43^+^ B cells and B1 cells in the bone marrow (Suppl. Fig. 1C). Compared with peritoneal macrophages, we found lower levels of both CD11b and SIRPα on B1 cells, clearly discriminating B1 cells from myeloid cells (Fig. 1D). Expression of SIRPα mRNA was confirmed by qRT-PCR on FACS sorted peritoneal B1a cells (CD19^+^CD5^+^CD11b^+^) (Figure 1E).

**Figure 1.**
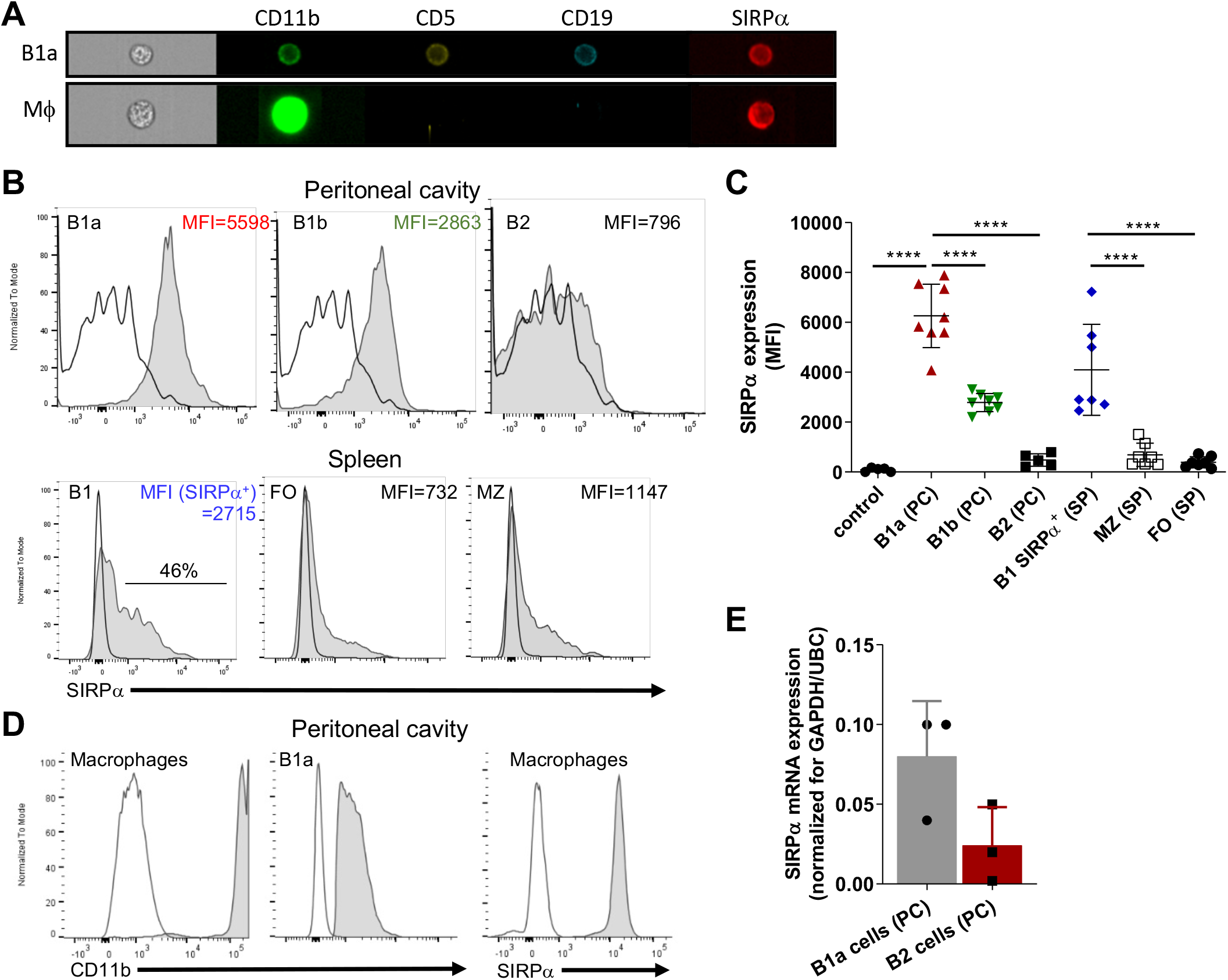
SIRPα is expressed by B1 cells. **A,** Imaging flow cytometry visualizing expression of SIRPα on CD19+CD5+CD11b+ B1a cells and CD11b+CD19-macrophages. Representative histograms (**B**) and synopsis (**C**) of SIRPα surface expression on defined B cell subpopulations as determined by flow cytometry (MFI, Mean fluorescence intensity). Note that the most prominent expression occurs on peritoneal cavity (PC) B1a and B1b cells, and on a subset of the splenic (SP) B1, with little or no expression on marginal zone (MZ) and follicular (FO) B cells. **D**, Macrophages cells from the peritoneal cavity show relatively high levels of expression of both CD11b and SIRPα. **E**, SIRPα mRNA expression on FACS sorted peritoneal cavity B1a and B2 cells. Data are in **C** and **E** are presented as mean±SEM and represent measurements of 5-8 and 3 individual mice, respectively. Statistical analysis was performed by one-way ANOVA with Dunnett’s multiple comparison corrrection, *p<0.05; ***p<0.001, ****p<0.0001.

### SIRPα limits natural IgM antibody levels in vivo

Because SIRPα, like Siglec-G, is also a typical inhibitory immunoreceptor with cytoplasmic ITIM motifs signaling through SHP-1 and/or SHP-2, we tested whether the lack of SIRPα signaling would affect natural antibody generation as well. As can be seen in Fig. 2A there was a prominent (~2-fold) enhancement in the plasma levels of total and OSE-reactive natural IgM antibodies in mice lacking the cytoplasmic tail of SIRPα (SIRPα^ΔCYT^ mice). This occurred for all OSE tested, including the so-called T15 epitope that defines PC-reactive EO6 type anti-OxLDL IgM antibodies with anti-atherogenic potential *in vivo* (Binder et al., 2003; Que et al., 2018), as well as phosphocholine (PC) and those against *ex vivo* oxidized LDL (i.e. MDA-LDL and Cu-OxLDL). Consistent with a specific B1 cell phenotype and a selective regulation of natural IgM levels the corresponding IgG levels were not altered (Fig. 2B). When SIRPα^ΔCYT^ mice were immunized with a typical T-cell independent type 2 (TI-2) antigen, DNP-Ficoll, we observed robust and enhanced DNP-specific IgM and IgG3 immune responses (Fig. 2C). These results indicate that SIRPα signaling regulates natural antibody production as well as the response of B1 cells to antigenic stimulation.

**Figure 2.**
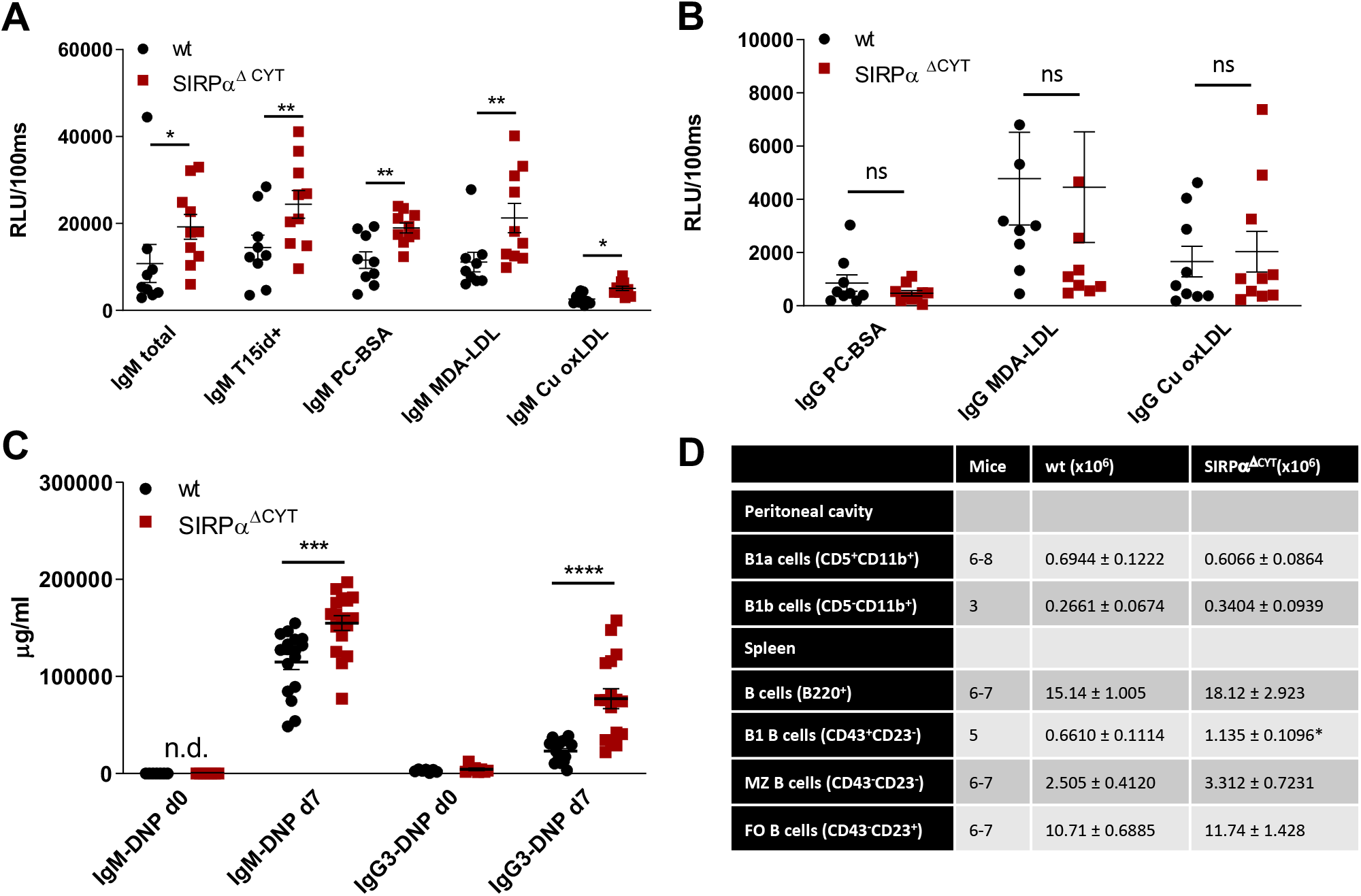
Loss of SIRPα signaling promotes B1 cell accumulation in the spleen and natural IgM antibody formation *in vivo*. Defective SIRPα signaling in mice lacking the SIRPα cytoplasmic tail (SIRPα^∆CYT^) results in increased plasma levels of natural IgM (**A**) but not IgG (**B**) antibodies directed against the indicated oxidation-specific epitopes under steady-state conditions; wt, wild-type mice. **C,** Immunization with the haptenated TI-2 antigen DNP-Ficoll triggers increased production of both IgM and IgG3 antibodies against DNP in SIRPα^∆CYT^ mice. Data are presented as mean ±SEM and are representative of 9-10 (**A**), 8 (**B**), 17 (**C)** individual mice. **D,** B cell numbers in peritoneal cavity and spleen of wt and SIRPα^∆CYT^ mice. Absolute number of different B cell populations was determined in the peritoneal cavity and the spleen of young adult mice (8-12 weeks) under steady state conditions. For further details see Suppl. Fig. 2A-G. Statistical analysis was performed by unpaired Student t-test, corrected for multiple comparisons with Holm-Sedak method where applicable, *p<0.05; **p<0.01, ***p<0.001, ****p<0.0001.; ns, non-significant; n.d., not detectable.

### SIRPα increases splenic B1 cell numbers without affecting B cell receptor function

Next, we investigated whether the changes in B1-cell associated antibodies in SIRPα^ΔCYT^ mice could be related to changes in B1 cell numbers in the peritoneal cavity or the spleen of these mice (Fig. 2D, Suppl. Fig. 2A-G). We did not detect significant differences in proportions or the absolute numbers of peritoneal cavity B1a and B1b cells. In contrast, proportions and absolute numbers of B1 cells in the spleen of SIRPα^ΔCYT^ mice were significantly (~2-fold) increased compared to wt animals. There were no significant differences in total cell numbers or proportions of other splenic B cell subsets, indicating that the effects were specific for B1 cells. We next asked how SIRPα might contribute to the increase in splenic B cell numbers and natural antibody levels. One possibility was that SIRPα was controlling the activation and expansion of B1 cells. First, we tested whether antigen recognition by B1 cells might be altered, but we observed no difference in e.g. the binding of a typical B1a cell antigen phosphatidylcholine (PtC) (Berland and Wortis, 2002) to B1a cells (Fig. 3A, Suppl. Fig. 2H). Next, we explored potential differences in B1a cell activation capacity. This included measuring Ca^2+^ flux upon cross-linking of BCR (Fig. 3B), and monitoring sorted PC B1a cell IgM secretion upon stimulation with LPS *in vitro* (Fig. 3C), a potent and typical B1 cell activation stimulus (Yang et al., 2007). Both read-outs showed comparable activation capacity of wt and SIRPα^ΔCYT^ B1a cells. Taken together, these findings support the idea that SIRPα signaling controls splenic B1 cell accumulation and, likewise as a consequence, also the levels of naturally occurring antibodies in a unique manner, but this was apparently not linked to direct regulation of B1 cell activation capacity.

**Figure 3.**
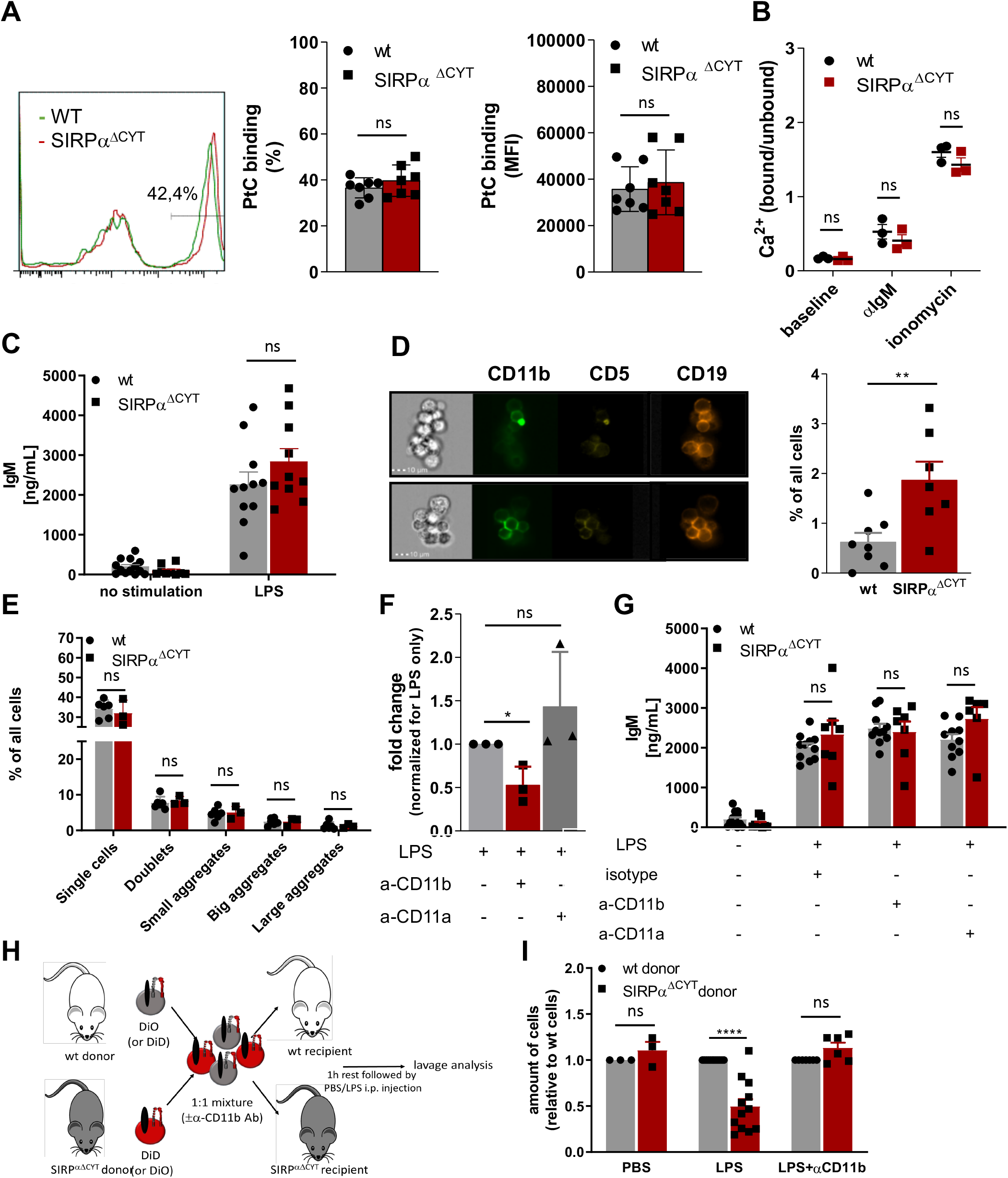
Loss of SIRPα signaling promotes LPS-induced CD11b/CD18 integrin-dependent B1 cell efflux from the peritoneal cavity. **A,** Comparable frequency (FACS plot of representative example, left panel; % positivity, middle panel) and magnitude (MFI, right panel) of phosphatidylcholine (PtC; a typical B1a antigen) antigen binding by wt and SIRPα^ΔCYT^ peritoneal cavity B1a cells (n=7 mice/group). **B**, Similar levels of B1a cell surface IgM signaling, as determined by intracellular Ca^2+^ mobilization, in wt and SIRPα^ΔCYT^ peritoneal cavity B1 cells triggered by anti-IgM antibodies; responses with ionomycin are shown as a positive control (n=3 mice/group). **C**, Comparable levels of LPS-stimulated IgM production in B1a cells isolated from SIRPα^ΔCYT^ and wt mice (n=10-11 mice/group). **D, E**, Lack of SIRPα signaling promotes the formation of large aggregates in isolated SIRPα^ΔCYT^ peritoneal cavity (**D**) but not splenic (**E**) B1a cells (n=6-8 mice/group). **F**, Enhanced frequency of B1 cell large aggregates in SIRPα^ΔCYT^ peritoneal cavity B1a cells relative to wt cells is CD11b/CD18-integrin-dependent as it is reduced by blocking anti-CD11b, but not anti-CD11a antibodies. **G**, Blockade of CD11b/CD18 or CD11a/CD18 integrins has no effect on IgM antibody production by peritoneal cavity B1 cells upon LPS stimulation. **H, I**, Experimental design of egress of adoptively transferred mixed SIRPα^ΔCYT^/wt B1 cells from the peritoneal cavity. **G**, Evaluation of adoptively transferred mixed SIRPα^ΔCYT^/wt B1 cells shows increased efflux of SIRPα^ΔCYT^ B1 cells relative to wt from the peritoneal cavity upon LPS stimulation and this enhanced egress is CD11b/CD18 dependent (n=3-12 mice/group). Similar data were obtained for SIRPα^ΔCYT^ recipients (not shown). Data are presented as mean ±SEM. Statistical analysis was performed by unpaired Student t-test, corrected for multiple comparisons with Holm-Sedak method where applicable, *p<0.05; **p<0.01, ***p<0.001, ****p<0.0001; ns, non-significant.

### SIRPα regulates CD11b/CD18 integrin function and B1 cell migratory capacity

Notably, when analyzing LPS-stimulated B1 cells by flow cytometry, we observed the presence of cell clusters in the cultures that appeared larger for SIRPα^ΔCYT^ B1a cells relative to their wt counterparts (Suppl. Fig. 4A). This prompted us to visualize and quantify this aggregate formation of B1a cells by imaging flow cytometry, which indeed demonstrated a substantially increased proportion of large aggregates (i.e. consisting of more than 4 cells) in SIRPα^ΔCYT^ peritoneal B1a cells as compared to wt B1a cells after LPS stimulation (Fig. 3D, Suppl. Fig. 4B,C). In contrary, we could not observe comparable aggregates when sorted splenic B1 cells were cultured in a similar manner (Fig. 3E). Interestingly, such doublets and large aggregates specific for CD11b^+^ B1 cells have been previously reported by Ghosn et al. and their formation seems to be dependent on CD11b (Ghosn et al., 2008). Furthermore, SIRPα inhibitory signaling has been previously linked to integrin function (Inagaki et al., 2000). We therefore hypothesized that SIRPα may serve as a negative regulator of CD11b/CD18 integrin function in B1 cells. To test this, we stimulated sorted peritoneal B1a cells with LPS in the presence of a blocking anti-CD11b antibody. Clearly, the increased formation of large aggregates triggered by LPS in mice lacking SIRPα could be partially prevented by blocking CD11b, but not by blocking CD11a (Fig. 3F). Next, we asked whether B1 cell aggregate formation through CD11b/CD18 integrin could be a prerequisite for production of natural antibodies. We tested supernatants of B1a cells that were activated by LPS in the presence of blocking CD11b antibody or blocking CD11a antibody. It appeared that SIRPα^ΔCYT^ B1a cells have comparable antibody production as wt B1 cells *in vitro*, independently of CD11b or CD11a function (Fig. 3G). Thus, whereas CD11b-mediated formation of large aggregates did not regulate natural antibody production *in vitro*, such B1 cell aggregate formation, which was promoted upon disruption of SIRPα signaling, nevertheless appeared a read-out for CD11b/CD18 activation. This suggested that SIRPα signaling was negatively regulating B1 cell integrin function. Of interest, Waffarn et al. have shown, that CD11b/CD18, unlike CD11a/CD18, is indispensable for migration of stimulated B1 cells from cavities to secondary lymphoid tissues where they mature into natural antibody producing plasma cells (Waffarn et al., 2015). This led us to propose that SIRPα could actually regulate CD11b/CD18 function during migration of B1 cells to the secondary lymphoid tissues, which would provide an explanation for the increased numbers of B1 cells in the spleens of SIRPα^ΔCYT^ mice. To directly test the effect of SIRPα signaling on the capacity of B1 cells to migrate from the peritoneal cavity we performed adoptive transfer experiments. To confirm integrin-dependence, B1 cells of both wt and SIRPα^ΔCYT^ donor mice were in parallel pre-incubated with blocking CD11b antibody, excessive amount of the antibody was washed away, and then the cells were labeled with unique membrane dyes, mixed in 1:1 ratio, and injected to either wt or SIRPα^ΔCYT^ recipients (Fig. 4H). This set-up allowed us to selectively monitor, in individual animals, the effect of SIRPα and CD11b/CD18 on the migration of B1 cells from the peritoneal cavity to the spleen. As expected, B1 cells that lack inhibitory cytoplasmic tail of SIRPα showed increased efflux from the peritoneal cavity (Fig. 4I). The enhanced exit of SIRPα^ΔCYT^ B1 cells was fully dependent on CD11b/CD18, as it could be completely inhibited by antibody blocking. Taken together, our data strongly suggest that CD11b/CD18 function in B1 cells is under negative control of SIRPα and that in absence of SIRPα signaling B1 cells have a higher propensity to leave the peritoneal cavity, thereby contributing to an accumulation of B1 cells in the spleen and an enhanced systemic production of natural antibodies.

**Figure 4.**
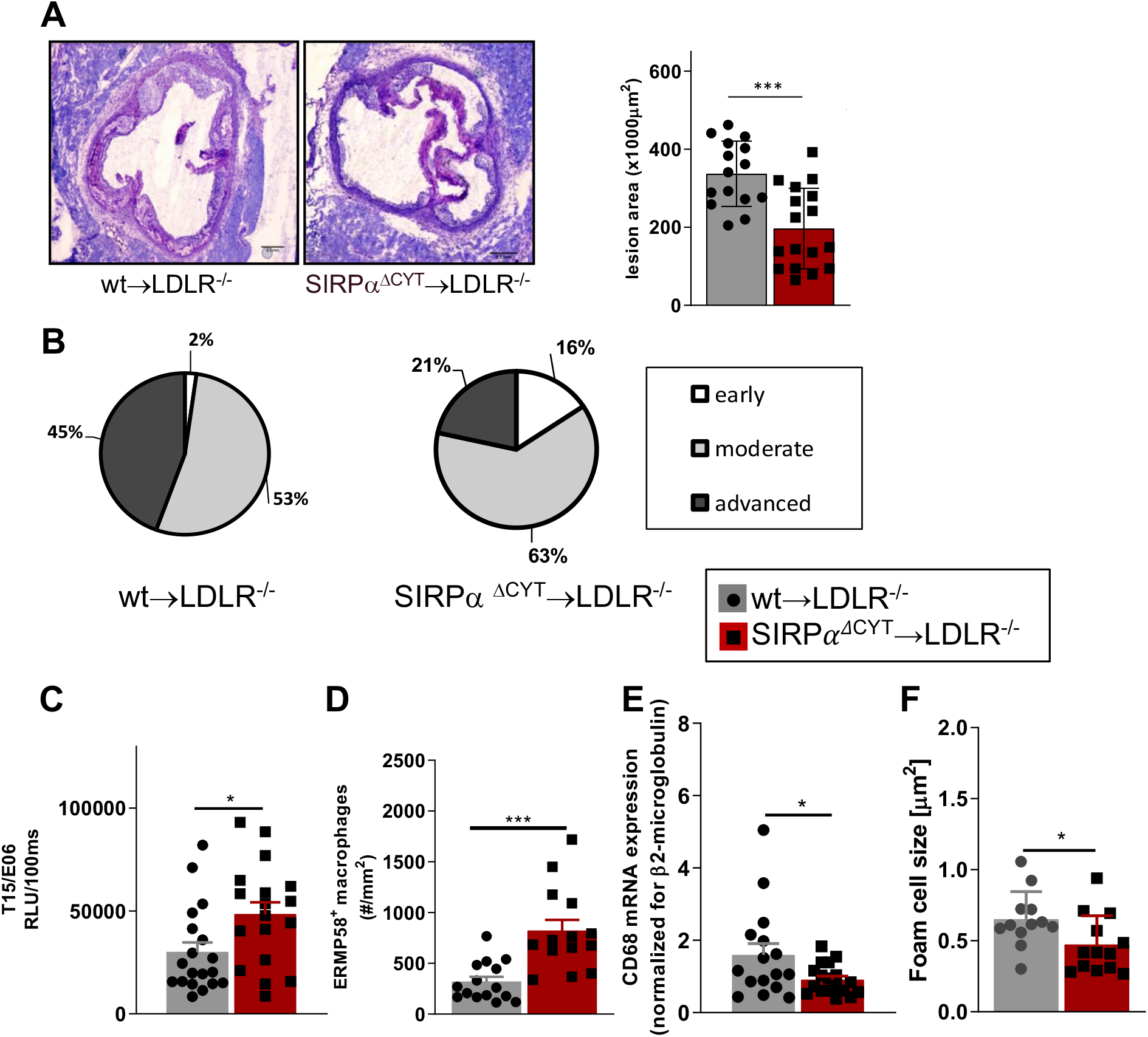
Loss of SIRPα signaling protects mice from atherosclerosis. **A, B**, Bone marrow chimeras carrying dysfunctional SIRPα in their hematopoietic compartment are protected from atherosclerosis, showing (**A**) smaller and (**B**) less severe aortic lesions. **C**, Atheroprotection in SIRPα^ΔCYT^>LDLR^−/−^ chimeras is associated with increased levels of oxLDL-targeting natural antibody T15/E06 in plasma. **D-F**, Atherosclerotic lesions of SIRPα^ΔCYT^>LDLR^−/−^ chimeras contain more small macrophages (**D**), less CD68+ macrophages (**E**), and a smaller foam cell area (**F**) as compared to wt chimeras. Data are presented as mean ±SEM and are representative of 15-17 (**A** and **B**), 18-19 (**C**), 12-16 (**D-F**) individual mice. Statistical analysis was performed by unpaired Student t-test used per variable, corrected for multiple comparisons with Holm-Sedak method, *p<0.05; **p<0.01, ***p<0.001, ****p<0.0001 or Chi-square test (**B**), *p<0.05.

### Lack of SIRPα signaling protects mice from atherosclerosis

In order to further establish the potential pathological/clinical relevance of the regulation of natural IgM antibody production by SIRPα *in vivo*, we decided to explore the role of SIRPα in atherosclerosis. Natural IgM antibodies produced by B1a cells have a well-established protective role in various diseases, and particularly in atherosclerosis, a feature which is apparently due to their capacity to neutralize oxLDL uptake and enhance apoptotic cell clearance by macrophages (Binder et al., 2003; Caligiuri et al., 2007; Faria-Neto et al., 2006). To directly address the role of SIRPα in atherosclerosis in mice, we transplanted wt and SIRPα^ΔCYT^ bone marrow into atherosclerosis-prone LDLR^−/−^ recipient mice. This well-established atherosclerosis model includes the replacement of peritoneal B cell populations (including B1 cells) by the donor cells (Binder et al., 2003; Caligiuri et al., 2007; Ding et al., 2007; Duber et al., 2009; Hoffmann et al., 2007; Holodick et al., 2009; Jellusova et al., 2010). Mice transplanted with wt or SIRPα^ΔCYT^ cells, and subjected to a high-fat diet, showed neither differences in weight, plasma cholesterol and triglyceride levels nor prominent changes in blood cell composition (Suppl. Fig. 3A-E). However, when the atherosclerotic lesions of these mice were evaluated, it became apparent that mice transplanted with SIRPα^ΔCYT^ cells developed much smaller lesions (Fig. 4A) with a substantially less severe phenotype (Fig. 4B) compared to wt chimeras. Additionally, when the plasma of atherosclerotic mice was analyzed for the presence of antibodies against oxLDL, in mice transplanted with SIRPα^ΔCYT^ cells elevated levels of T15/E06 IgM (Fig. 4C), an oxLDL neutralizing antibody, being particularly critical in the protection against atherosclerosis, were found. (Binder et al., 2004; Binder et al., 2003; Que et al., 2018). Similar to the steady state situation, levels of IgG targeting OSE remained unaltered (Suppl. Fig. 3F). A more detailed evaluation of the cellular composition of the lesions showed an increase in the number of newly recruited ERMP58+ myeloid cells (Fig. 4D), which was associated with a significantly decreased mRNA for CD68^+^ macrophages (Fig. 4E) and size of the area occupied by foam cells in mice transplanted with SIRPα^ΔCYT^ cells (Fig. 4F), consistent with the proposed mechanism of IgM-mediated inhibition of foam cell formation (Binder et al., 2003; Feng et al., 2010; Horkko et al., 1999). There were no significant differences observed in numbers of other immune cells known to be involved in pathogenesis of atherosclerosis, such as T-lymphocytes and neutrophils (Suppl. Fig. 3G,H).

## Discussion

In this study we demonstrate the expression and functional relevance of the inhibitory receptor SIRPα on the B1 cells in mice. Our results show that SIRPα is an inhibitory receptor on B1 cells controlling the numbers of splenic B1 cells, thereby most likely affecting systemic natural antibody levels. The increase in splenic B1 cell numbers in the absence of SIRPα occurs most probably as a consequence of its missing inhibitory effect on CD11b/CD18 integrin, which promotes the migration of these B1 cells from the peritoneal cavity to the spleen. We also show that lack of inhibitory SIRPα signaling in atherosclerotic mice leads to selectively elevated plasma levels of oxLDL-neutralizing natural antibodies and propose that this directly contributes to the atheroprotective effect of SIRPα deletion. The latter is in agreement with the well-established regulatory role of such antibodies in atherosclerosis (Binder et al., 2016).

SIRPα is one of the most abundant inhibitory receptors on myeloid cells including neutrophils, monocytes, the majority of tissue macrophages and CD4^+^ dendritic cells, affecting a variety of cell functions in a primarily negative fashion (Barclay and van den Berg, 2014; Matlung et al., 2017). SIRPα has, as yet, not been reported to be expressed by any B cells. Recently, SIRPα was also reported to be selectively expressed on a small subset of T lymphocytes, i.e. exhausted CD8+ memory T cells emerging after chronic viral infection (Myers et al., 2019). For a long time the general assumption has been that SIRPα is, at least among hematopoietic cells, restricted to the myeloid lineage (Adams et al., 1998; Brooke et al., 1998). Most of the studies based the absence of SIRPα from lymphocytes on staining of blood cells in rodents (Adams et al., 1998; Sato-Hashimoto et al., 2011) while other, less accessible or more obscure subpopulations of lymphoid origin have remained unexplored. We found expression of SIRPα exclusively on B1 cells in the peritoneal cavity and on a minor subset of B cells in the spleen. We also observed that the population of steady-state splenic B1 cells is roughly doubled in mice lacking SIRPα signaling. This increase in splenic B1 cells may well explain the (also ~2-fold) higher IgM plasma levels found in SIRPα^ΔCYT^ mice. Antigens with repetitive patterns, such as lipids and glycolipids, incl. self-antigens that are generated e.g. upon oxidation or apoptosis can induce multivalent antigen crosslinking of specific BCR on B1 cells and induce TI-2 responses. Due to this self- and poly- reactivity of B1 cells, their functions have to be tightly regulated to avoid autoimmunity and several inhibitory receptors with diverse regulatory functions on B1 cells are known to be instrumental in this. We have observed that lack of SIRPα on B1 cells has no measurable effect on calcium flux and IgM secretion, which may suggest that Siglec-G and CD5, which have previously shown to regulate these parameters, comprise the major regulators of BCR signaling in B1 cells (Bikah et al., 1996). SIRPα^ΔCYT^ mice have moderately increased numbers of B1 cells only in the spleen with the peritoneal population virtually unaltered. Also, B1-associated antibody levels are elevated in SIRPα^ΔCYT^ mice, both at baseline and after TI-2 antigen exposure. Natural antibodies are predominantly produced by B1 cells in secondary lymphoid organs such as spleen (Baumgarth, 2011) or specific (e.g. mediastinal) lymph nodes (Waffarn et al., 2015). Egress of B1 cells residing in the body cavities (peritoneal or pleural) into the secondary lymphoid organs has been demonstrated to depend on the CD11b/CD18 integrin (Waffarn et al., 2015). CD11b can function only in heterodimer with CD18 integrin, forming together CD11b/CD18 (also known as CR3, Mac-1, Integrin alpha M, or α_M_β_2_ integrin). B1 cells are known to express various integrin molecules, but B1 cells are the only B cells that express CD11b/CD18 integrin and until now no direct regulator of its function has been described. We show here that SIRPα negatively regulates capacity of B1 cells to exit the peritoneal cavity through CD11b/CD18 integrin, identifying this inhibitory immunoreceptor as the first *bona fide* regulator of B1 CD11b/CD18 function and migration. Therefore, it seems that B1 cells are subject to extensive and versatile control via immune checkpoints. In fact, a picture emerges where different B1 cell functions appear to be regulated by distinct inhibitory receptors, with SIRPα controlling their migratory behavior, whereas others e.g. Siglec-G and CD5 directly control their activation.

The homeostatic function of IgM antibodies has been well documented in atherosclerosis (Binder et al., 2003; Caligiuri et al., 2007; Faria-Neto et al., 2006; Gronwall and Silverman, 2014). Mice lacking SIRPα signaling in the hematopoietic compartment showed increased plasma levels of T15/E06 IgM and show smaller and less severe atherosclerotic lesions. This is similar to observations with Siglec G bone marrow chimeras where lack of inhibitory signaling by Siglec G led to increased OSE-specific natural IgM antibody levels and decreased atherosclerosis development (Gruber et al., 2016). Increased plasma level of T15/E06 IgM is a very likely mechanism of atheroprotection in SIRPα^ΔCYT^ chimeras. However, transplantation of SIRPα^ΔCYT^ and wt bone marrow resulted in replacement of both myeloid and lymphoid lineage in the donor LDLR^−/−^ mice. As macrophages are important players in development of atherosclerosis we cannot exclude their contribution to the observed phenotype, especially since the SIRPα counter-receptor CD47 appears involved in pathogenesis of atherosclerosis through inhibition e.g. macrophage efferocytosis (Kojima et al., 2016). Whether antibodies targeting SIRPα, rather than CD47, would show similar effect still remains to be confirmed, also because the CD47 monoclonal antibody miap410 used in the study of Kojima (Kojima et al., 2016) has prominent opsonizing capacity and a less convincing ability to actually block CD47-SIRPα axis (Han et al., 2000). Of relevance, population studies have shown that increased levels of antibodies against OSE are correlated with better prognosis in cardiovascular diseases, suggesting the existence of similar natural antibody producing B cells in humans (Ravandi et al., 2011).

Collectively, our data identify SIRPα as a novel B1 cell immune checkpoint, which functions to control B1 cell migration, and imply SIRPα as a potential therapeutic target in atherosclerosis. The CD47-SIRPα innate immune checkpoint is currently extensively studied in the context of cancer immunotherapy (Matlung et al., 2017; van den Berg and Valerius, 2019), with a number of different agents in preclinical and/or clinical development, and ~20 ongoing clinical trials, carving out a path for potential therapeutic targeting of the CD47-SIRPα axis also in other diseases, including cardiovascular disease.

## Materials and Methods

### Mice

C57BL/6 mice with a targeted deletion of the SIRPα cytoplasmic region have been described previously (Inagaki et al., 2000; Yamao et al., 2002). The mice that were originally generated onto the 129/Sv background were backcrossed onto C57BL/6 mice for at least 13 generations. Wild-type (wt) C57BL/6 mice of the same genetic background were maintained under specific pathogen-free conditions together with the SIRPα^ΔCYT^ mice in the breeding facility of the Netherlands Cancer Institute, Amsterdam, the Netherlands or the VU Medical Center, Amsterdam, the Netherlands. Bone marrow was isolated and used for transplantation at the animal facility of Maastricht University, Maastricht, the Netherlands. Animals were housed in ventilated cages and treated according to European Commission guidelines. They were euthanized using combination of isofluran and CO_2_. All animal experiments were approved by the Animal Welfare Committee of the VU Medical Center Amsterdam, the Netherlands, Maastricht University, Maastricht, the Netherlands and The Netherlands Cancer Institute, Amsterdam, the Netherlands. LDLR^−/−^ mice on C57BL/6J background were obtained from Jackson Laboratory (Bar Harbor, ME, USA).

### Flow cytometry analysis of SIRPα expression on mouse B cells

Mouse B cells were isolated from the peritoneal cavity by peritoneal lavage of 8-12 weeks old SIRPα^ΔCYT^ mice and age matched wt mice. Mice were sacrificed and immediately after that 5 mL of cold PBS containing 3% of fetal calf serum (FCS) and 3mM EDTA was injected into their peritoneal cavity. After gentle massage, cells were collected and used for analysis of SIRPα expression. Additionally, bone marrow and spleens of the same mice were isolated and blood samples were taken to analyze for expression of SIRPα. Fetal livers were isolated from mice of FVB background at E12. Single cell suspensions of splenocytes were prepared after the spleens were homogenized through 100μm filter (BD Biosciences, Bedford, MA, USA), lysed with lysis buffer and washed twice with cold PBS. For blood analysis whole blood was first spun down at 2000 rpm at 4°C for 10 min and plasma was collected and stored at −80°C for later analysis of antibodies level. Erythrocytes were lysed using cold lysis buffer containing 155mM NH_4_Cl, 10mM KHCO_3_ and 0.1mM EDTA (ethylene diamine tetra acetic acid), pH 7.4. For flow cytometry analysis first Fc receptors were blocked using α-CD16/CD32 antibody (clone 2.4G2, BD Biosciences, Bedford, MA, USA). The cells were subsequently washed and stained for following surface markers with directly conjugated antibodies against CD19/B220 (PerCP Cy5.5 or eFluor 450), CD11b (Alexa Fluor 488), CD5 (PE), IgM (PE), SIRPα (APC or PerCP 710) (all antibodies purchased from eBioscience, San Diego, CA, USA) and CD43 (APC Cy7, BioLegend, San Diego, CA, USA). Expression of proteins was measured using FACS Canto II HTS (BD Biosciences, Bedford, MA, USA) and analyzed using FlowJo software (FlowJo LLC, Ashland, OR, USA). To reliably detect SIRPα expression and separate it from autofluorescence, fluorescence minus one (FMO) control was applied, when cells were stained for all determinants except SIRPα (Ghosn et al., 2008).

### Quantitative RT-PCR to determine SIRPα mRNA expression

RNA was isolated from FACS sorted mouse B1a and B2 cells based on markers listed above with QIAamp RNA Blood mini kit according to manufacturer’s instructions (Qiagen, Venlo, The Netherlands). RNA was eluted with 30μL H2O, to obtain as high as possible concentration of RNA. Total RNA was reverse transcribed using the III first-strand synthesis system for RT-PCR (Invitrogen, Breda, the Netherlands). In short, 8 μL RNA was primed with 2,5 μM oligo-dT primer which specifically targets mRNA and 0,5 mM dNTP for 5 minutes at 65°C. Reverse transcription was performed with 10 U/μL Superscript III in the presence of 5 mM MgCl_2_, 20 mM Tris-HCL, and 50 mM KCl, pH8,4 (RT buffer), 2 U/ μL RNAseOUT™, lacking DTT for reasons described before(Lekanne Deprez et al., 2002) for 50 minutes at 50°C. After that,

Superscript III was inactivated by incubation for 5 minutes at 85°C, followed by chilling on ice. Immediately thereafter, 2 U RNase H was added and incubated at 37°C for 20 minutes. Subsequently cDNA was stored at −20°C until further use. Amplification by PCR was performed on a LightCycler instrument (Roche, Almere, the Netherlands), with software version 3.5. The reaction was performed with Lightcycler FastStart DNA Master^PLUS^ SYBR Green I (Roche, Almere, the Netherlands). The annealing temperature used for all primers was 60°C. The reaction mix consisted of 4 μL of cDNA, 1 μM of each primer combination and 4 μL of Lightcycler FastStart DNA Master^PLUS^ SYBR Green I (Roche) in a total volume of 20 μL. After an incubation step for 10 minutes at 95°C, the template was amplified for 45 cycles at 95°C, annealing of the primers was performed at 60°C for 30 seconds, followed by extension at 72°C for 15 seconds. At the end of the 45 cycles, a melting curve was generated to determine the unique features of the DNA amplified. cDNA of control wt animals was used as a standard curve with a serial 10-fold dilution. Musculus Ubiquitin C was used as a reference gene. The product was sequenced by Big-dye Terminator Sequencing and ABI Prism software (Applied Biosystems, Foster City, USA). The sequence obtained was verified with BLAST (http://www.ncbi.nlm.nih.gov/BLAST/) to determine specificity. Primer sequences are available upon request.

### Binding of phosphatidylcholine by primary mouse B cells

B cells were isolated and labeled with antibodies against surface CD5 and B220 as described above along with fluorescein-labeled phosphatidylcholine (PtC) liposomes (DOPC/CHOL 55:45, Formumax Scientific Inc.). The cells were incubated on ice for 20 min followed by two washing steps, then cells were analyzed using an LSRII flow cytometer (BD Biosciences, Bedford, MA, USA) for binding of phosphatidylcholine and data were processed with FlowJo software (FlowJo LLC, Ashland, OR, USA).

### Intracellular calcium mobilization measurement in primary mouse B1a cells

B cells were isolated from the peritoneal cavity of 8-12 weeks old SIRPα^ΔCYT^ mice and aged matched wt mice using peritoneal lavage as described above. First, Fc receptors were blocked using α-CD16/CD32 antibody (clone 2.4G2, BD Biosciences, Bedford, MA, USA). The cells were then stained with directly labeled antibody against CD5 (APC) and B220 (APC Cy7, both

BD Biosciences, Bedford, MA, USA) allowing identification of B1a cells. Calcium flux was determined as described before by flow cytometric determination (Muggen et al., 2015). BCR-mediated Ca2^+^ mobilization was measured for 60s after the cells were stimulated either with 10 μg/mL F(ab′)2 of polyclonal goat anti-mouse IgM (Jackson ImmunoResearch, West Grove, PA, USA) or 0.5mM phosphatidylcholine (PtC) (F60103F-F, FormuMax USA). At the end of each Ca^2+^ measurement cells were treated with ionomycine (Life Technologies, Eugene, OR, USA) as a positive control for calcium signaling. Data were acquired on an LSRII flow cytometer (BD Biosciences, Bedford, MA, USA) and data analysis was performed with the use of FlowJo software (FlowJo LLC, Ashland, OR, USA).

### Immunization of mice with DNP-Ficoll

For B1-specific immunization intraperitoneal injection of TI-2 antigen di-nitro phenyl (DNP)-Ficoll was used as described before (Maas et al., 1997). Mice were injected with 50μg of DNP-Ficoll in 200μL PBS solution or with 200μL of PBS only as control. After 7 days animals were sacrificed and their blood was collected, plasma was harvested and stored at −80°C before analysis of IgM and IgG3 antibodies by ELISA.

### Measurement of IgM and IgG with ELISA

Plasma levels of IgM antibodies against several OSE were determined as described before (Chou et al., 2009). Dilutions of 1:100 (anti-phosphocholine (PC)-BSA IgM and all IgG antibodies), 1:500 (E06/T15id+ IgM, anti-malondialdehyde (MDA-)LDL IgM, anti-Cu-OxLDL IgM) and 1:20.000 (total IgM) were used. Supernatants of peritoneal B1 cell cultures or plasma of mice were serially diluted to determine IgM production after 48h of stimulation or IgM/IgG3 against DNP-Ficoll after 7-days immunization as previously described (Ha et al., 2006). Briefly, supernatants were measured by sandwich ELISA, using unlabeled for coating and peroxidase-labeled anti-mouse IgM/IgG antibody (total, or DNP-Ficoll specific, Southern Biotechnology, Birmingham, AL, USA) for detection, and azino-bis-ethylbenz-thiazoline sulfonic acid was used as the substrate. Antibody concentrations were calculated by using purified mouse IgM protein (IgM DNP-Ficoll and IgG3 DNP-Ficoll PMP52, Serotec, UK) as a standard.

### Stimulation of peritoneal B1 cell

Peritoneal B1a cells were obtained through negative magnetic-activated cell separation with a cocktail of antibodies depleting other than B1a cells achieving more than 90% purity in isolation (Miltenyi Biotec B.V., Utrecht, The Netherlands). B1a cells were subsequently counted and plated in IMDM medium (Invitrogen, Eugene, OR, USA) supplemented with 10% fetal calf-serum (FCS; Bodinco, Alkmaar, The Netherlands, 100 U/mL of penicillin, 100 mg/mL of streptomycin, and 2 mM L-glutamine (all Gibco Invitrogen, Breda, The Netherlands), and beta-mercapthoethanol (3.57 × 10^−4^M; Millipore, Amsterdam, the Netherlands). Cells were plated in 96-well plate in density of 1×10^6^/mL in 200μL of medium and cultured at 37°C and 5% CO2 for 48h in presence of 5μg of lipopolysaccharide (LPS, isolated from E. coli strain 055:B5, Sigma, St. Louis, MO, USA; LBP from R&D Systems, Abingdon, UK), isotype control (rat IgG2b, eBioscience, San Diego, CA, USA), anti-CD11b antibody (functional grade, eBioscience, San Diego, CA, USA), or anti-CD11a (functional grade, eBioscience, San Diego, CA, USA) in final concentration 10μg/mL. After that supernatant was collected and stored at −80°C before measurement of IgM antibody by ELISA. Cells were harvested and processed for analysis by flow cytometry and imaging cytometry.

### Image Stream analysis of aggregates formation

LPS-stimulated B1a cells were stained with following antibodies: CD19 (PerCP Cy5.5), CD11b (Alexa Fluor 488), CD5 (PE), SIRPα (APC), (all antibodies purchased from eBioscience, San Diego, CA, USA) and analyzed imaging cytometry to detect formation of aggregates (Image Stream, (Image Stream, Amnis, EMD, Millipore, Seattle, WA, USA) with gating strategy as follows: all events were divided based on their size into single cells (1 cell); doublets (2 cells); doublets and small aggregates (2-3 cells); big aggregates (3-4 cells); and large aggregates (>4 cells). Analysis of data was performed using analysis software IDEAS (Amnis Corporation, Seattle, WA, USA) and depicted as percentage of all gated events.

### Adoptive transfer of peritoneal B1 cells

Either wt or SIRPα^ΔCYT^ animals were used as donors of peritoneal B1 cells for adoptive transfer. Cells were harvested by peritoneal lavage as described above. The cells were left in IMDM medium supplemented with 10% FCS (Bodinco, Alkmaar, The Netherlands, 100 U/mL of penicillin, 100 mg/mL of streptomycin, and 2 mM L-glutamine (all Gibco Invitrogen, Breda, The

Netherlands) to rest for 30min. After that the easy to detach and floating cells (excluding adherent peritoneal macrophages) were first incubated with Fc receptor blocking antibody (anti-CD16/CD32) and after washing incubated with either isotype control (rat IgG2b, eBioscience, San Diego, CA, USA) or anti-CD11b antibody (functional grade, eBioscience, San Diego, CA, USA) in concentration of 10μg/mL for 30min. Antibodies were washed away and wt and SIRPα^ΔCYT^ cells were labeled with membrane dye DiD and DiO, respectively (or vice versa, to exclude effect of the dye on cell properties). Cells were washed multiple times and mixed in 50:50 ratio based on a cell count. The actual ratio was additionally determined by analyzing a small sample of pooled cells on flow cytometer allowing later normalization of the cell input. Cells were then injected into the peritoneal cavity of either wt or SIRPα^ΔCYT^ recipient mice, left resting for 1h, and followed by either 200μL injection of PBS (control) or 10μg of LPS in 200μL PBS intraperitoneally to induce migration of B1 cells from the peritoneal cavity (Ha et al., 2006; Yang et al., 2007). Peritoneal lavage of recipient mice was performed 3h after PBS/LPS injection. Lavage composition was analyzed by flow cytometry in CD19+ single cell population and percentage of cells with distinct membrane label was calculated. Percentage of cells was normalized for input of pooled cells as indicated above.

### Bone marrow transplantation

One week before transplantation, female LDLR^−/−^ mice were housed in filter top cages with neomycin (100 mg/L; Gibco, Breda, the Netherlands) and polymyxin B sulphate (66104 U/L; Gibco Breda, the Netherlands) in their acidified drinking water. The animals received 6 Gy of total body irradiation twice on consecutive days. Bone marrow isolated from SIRPα^ΔCYT^ and wt mice was injected intravenously to rescue the hematopoietic system of the irradiated mice as described previously (Goossens et al., 2010). Four weeks after the transplantation, mice were put on a high fat diet (0.15% cholesterol, 16% fat, Arie Blok, the Netherlands) for 10 weeks and level of chimerism was tested (reached 98.76%±0.73).

### Mouse blood parameters

Blood was withdrawn at the indicated times during high fat diet period and plasma lipid levels were enzymatically measured using ELISA (Sigma Aldrich, Zwijndrecht, the Netherlands).

### Atherosclerotic lesions analysis

Transplanted animals were sacrificed and isolated hearts were cut perpendicularly to heart axis just below the atrial tips, as described before (Kanters et al., 2004; Kanters et al., 2003). Serial cross sections were stained with toluidin blue and lesion areas were quantified using Adobe Photoshop software. Severity of lesions was scored as early, moderate and advanced, using criteria as described before (Kanters et al., 2004; Kanters et al., 2003). In detail, early lesions were fatty streaks containing only foam cells; moderate (intermediate) lesions were characterized by the additional presence of a collagenous cap, and advanced lesions showed involvement of the media mostly accompanied by increased collagen content and necrosis of the plaque. Foam cell size within plaques was determined by dividing the size of an allocated foamy macrophage area by the number of macrophages.

### Immunohistochemical staining

Atherosclerotic lesions from aortic roots were stained with various antibodies to identify neutrophils (NIMP directed against Ly6G, a gift from P. Heeringa), T cells (KT3, directed against CD3) and newly recruited macrophages (ER-MP58, a gift from P. Leenen) followed by detection with biotin labeled rabbit anti-rat antibody and staining with ABC kit (Vector Labs, Burlingame, CA).

### Statistical analysis

Statistical analysis was performed using GraphPad Prism version 8.02 (GraphPad Software, San Diego, CA, USA). Data were evaluated by two-tailed student t-test if two columns were compared. If more columns were compared, one-way ANOVA followed by multiple comparison correction was applied.

## Acknowledgements

The authors thank Martijn A. Nolte and Rene A.W. van Lier for useful discussions. This work was supported by NWO-TOP grant (#91208001) awarded to GK, MPJdW and TKvdB.

## References

Adams, S., L.J. van der Laan, E. Vernon-Wilson, L.C. Renardel de, E.A. Dopp, C.D. Dijkstra, D.L. Simmons, and T.K. van den Berg. 1998. Signal-regulatory protein is selectively expressed by myeloid and neuronal cells. J. Immunol 161:1853–1859.

Advani, R., I. Flinn, L. Popplewell, A. Forero, N.L. Bartlett, N. Ghosh, J. Kline, M. Roschewski, A. LaCasce, G.P. Collins, T. Tran, J. Lynn, J.Y. Chen, J.P. Volkmer, B. Agoram, J. Huang, R. Majeti, I.L. Weissman, C.H. Takimoto, M.P. Chao, and S.M. Smith. 2018. CD47 Blockade by Hu5F9-G4 and Rituximab in Non-Hodgkin’s Lymphoma. N Engl J Med 379:1711–1721.

Amezcua Vesely, M.C., M. Schwartz, D.A. Bermejo, C.L. Montes, K.M. Cautivo, A.M. Kalergis, D.J. Rawlings, E.V. Acosta-Rodriguez, and A. Gruppi. 2012. FcgammaRIIb and BAFF differentially regulate peritoneal B1 cell survival. J. Immunol 188:4792–4800.

Bagchi-Chakraborty, J., A. Francis, T. Bray, L. Masters, D. Tsiantoulas, M. Nus, J. Harrison, M. Broekhuizen, J. Leggat, M.R. Clatworthy, M. Espeli, K.G.C. Smith, C.J. Binder, Z. Mallat, and A.P. Sage. 2019. B Cell Fcgamma Receptor IIb Modulates Atherosclerosis in Male and Female Mice by Controlling Adaptive Germinal Center and Innate B1-Cell Responses. Arterioscler Thromb Vasc Biol ATVBAHA118312272.

Barclay, A.N., and T.K. van den Berg. 2014. The interaction between signal regulatory protein alpha (SIRPalpha) and CD47: structure, function, and therapeutic target. Annu. Rev. Immunol 32:25–50.

Baumgarth, N. 2011. The double life of a B-1 cell: self-reactivity selects for protective effector functions. Nat. Rev. Immunol 11:34–46.

Berland, R., and H.H. Wortis. 2002. Origins and functions of B-1 cells with notes on the role of CD5. Annu. Rev. Immunol 20:253–300.

Bikah, G., J. Carey, J.R. Ciallella, A. Tarakhovsky, and S. Bondada. 1996. CD5-mediated negative regulation of antigen receptor-induced growth signals in B-1 B cells. Science 274:1906–1909.

Binder, C.J., K. Hartvigsen, M.K. Chang, M. Miller, D. Broide, W. Palinski, L.K. Curtiss, M. Corr, and J.L. Witztum. 2004. IL-5 links adaptive and natural immunity specific for epitopes of oxidized LDL and protects from atherosclerosis. J. Clin. Invest 114:427–437.

Binder, C.J., S. Horkko, A. Dewan, M.K. Chang, E.P. Kieu, C.S. Goodyear, P.X. Shaw, W. Palinski, J.L. Witztum, and G.J. Silverman. 2003. Pneumococcal vaccination decreases atherosclerotic lesion formation: molecular mimicry between Streptococcus pneumoniae and oxidized LDL. Nat. Med 9:736–743.

Binder, C.J., N. Papac-Milicevic, and J.L. Witztum. 2016. Innate sensing of oxidation-specific epitopes in health and disease. Nat Rev Immunol 16:485–497.

Brooke, G.P., K.R. Parsons, and C.J. Howard. 1998. Cloning of two members of the SIRP alpha family of protein tyrosine phosphatase binding proteins in cattle that are expressed on monocytes and a subpopulation of dendritic cells and which mediate binding to CD4 T cells. Eur. J. Immunol 28:1–11.

Caligiuri, G., J. Khallou-Laschet, M. Vandaele, A.T. Gaston, S. Delignat, C. Mandet, H.V. Kohler, S.V. Kaveri, and A. Nicoletti. 2007. Phosphorylcholine-targeting immunization reduces atherosclerosis. J. Am. Coll. Cardiol 50:540–546.

Choi, Y.S., J.A. Dieter, K. Rothaeusler, Z. Luo, and N. Baumgarth. 2012. B-1 cells in the bone marrow are a significant source of natural IgM. Eur. J. Immunol 42:120–129.

Chou, M.Y., L. Fogelstrand, K. Hartvigsen, L.F. Hansen, D. Woelkers, P.X. Shaw, J. Choi, T. Perkmann, F. Backhed, Y.I. Miller, S. Horkko, M. Corr, J.L. Witztum, and C.J. Binder. 2009. Oxidation-specific epitopes are dominant targets of innate natural antibodies in mice and humans. J. Clin. Invest 119:1335–1349.

Ding, C., Y. Liu, Y. Wang, B.K. Park, C.Y. Wang, P. Zheng, and Y. Liu. 2007. Siglecg limits the size of B1a B cell lineage by down-regulating NFkappaB activation. PLoS. One 2:e997.

Duber, S., M. Hafner, M. Krey, S. Lienenklaus, B. Roy, E. Hobeika, M. Reth, T. Buch, A. Waisman, K. Kretschmer, and S. Weiss. 2009. Induction of B-cell development in adult mice reveals the ability of bone marrow to produce B-1a cells. Blood 114:4960–4967.

Faria-Neto, J.R., K.Y. Chyu, X. Li, P.C. Dimayuga, C. Ferreira, J. Yano, B. Cercek, and P.K. Shah. 2006. Passive immunization with monoclonal IgM antibodies against phosphorylcholine reduces accelerated vein graft atherosclerosis in apolipoprotein E-null mice. Atherosclerosis 189:83–90.

Feng, X., Y. Zhang, R. Xu, X. Xie, L. Tao, H. Gao, Y. Gao, Z. He, and H. Wang. 2010. Lipopolysaccharide up-regulates the expression of Fcalpha/mu receptor and promotes the binding of oxidized low-density lipoprotein and its IgM antibody complex to activated human macrophages. Atherosclerosis 208:396–405.

Ghosn, E.E., Y. Yang, J. Tung, L.A. Herzenberg, and L.A. Herzenberg. 2008. CD11b expression distinguishes sequential stages of peritoneal B-1 development. Proc Natl Acad Sci U S A 105:5195–5200.

Goossens, P., M.J. Gijbels, A. Zernecke, W. Eijgelaar, M.N. Vergouwe, d.M. van, I, J. Vanderlocht, L. Beckers, W.A. Buurman, M.J. Daemen, U. Kalinke, C. Weber, E. Lutgens, and M.P. de Winther. 2010. Myeloid type I interferon signaling promotes atherosclerosis by stimulating macrophage recruitment to lesions. Cell Metab 12:142–153.

Grasset, E.K., A. Duhlin, H.E. Agardh, O. Ovchinnikova, T. Hagglof, M.N. Forsell, G. Paulsson-Berne, G.K. Hansson, D.F. Ketelhuth, and M.C. Karlsson. 2015. Sterile inflammation in the spleen during atherosclerosis provides oxidation-specific epitopes that induce a protective B-cell response. Proc Natl Acad Sci U S A 112:E2030–2038.

Gronwall, C., and G.J. Silverman. 2014. Natural IgM: beneficial autoantibodies for the control of inflammatory and autoimmune disease. J. Clin. Immunol 34 Suppl 1:S12–S21.

Gruber, S., T. Hendrikx, D. Tsiantoulas, M. Ozsvar-Kozma, L. Goderle, Z. Mallat, J.L. Witztum, R. Shiri-Sverdlov, L. Nitschke, and C.J. Binder. 2016. Sialic Acid-Binding Immunoglobulin-like Lectin G Promotes Atherosclerosis and Liver Inflammation by Suppressing the Protective Functions of B-1 Cells. Cell Rep 14:2348–2361.

Ha, S.A., M. Tsuji, K. Suzuki, B. Meek, N. Yasuda, T. Kaisho, and S. Fagarasan. 2006. Regulation of B1 cell migration by signals through Toll-like receptors. J Exp Med 203:2541–2550.

Han, X., H. Sterling, Y. Chen, C. Saginario, E.J. Brown, W.A. Frazier, F.P. Lindberg, and A. Vignery. 2000. CD47, a ligand for the macrophage fusion receptor, participates in macrophage multinucleation. J Biol Chem 275:37984–37992.

Hayakawa, K., R.R. Hardy, D.R. Parks, and L.A. Herzenberg. 1983. The “Ly-1 B” cell subpopulation in normal immunodefective, and autoimmune mice. J Exp Med 157:202–218.

Hoffmann, A., S. Kerr, J. Jellusova, J. Zhang, F. Weisel, U. Wellmann, T.H. Winkler, B. Kneitz, P.R. Crocker, and L. Nitschke. 2007. Siglec-G is a B1 cell-inhibitory receptor that controls expansion and calcium signaling of the B1 cell population. Nat. Immunol 8:695–704.

Holodick, N.E., K. Repetny, X. Zhong, and T.L. Rothstein. 2009. Adult BM generates CD5+ B1 cells containing abundant N-region additions. Eur. J. Immunol 39:2383–2394.

Horkko, S., D.A. Bird, E. Miller, H. Itabe, N. Leitinger, G. Subbanagounder, J.A. Berliner, P. Friedman, E.A. Dennis, L.K. Curtiss, W. Palinski, and J.L. Witztum. 1999. Monoclonal autoantibodies specific for oxidized phospholipids or oxidized phospholipid-protein adducts inhibit macrophage uptake of oxidized low-density lipoproteins. J. Clin. Invest 103:117–128.

Inagaki, K., T. Yamao, T. Noguchi, T. Matozaki, K. Fukunaga, T. Takada, T. Hosooka, S. Akira, and M. Kasuga. 2000. SHPS-1 regulates integrin-mediated cytoskeletal reorganization and cell motility. EMBO J 19:6721–6731.

Jellusova, J., S. Duber, E. Guckel, C.J. Binder, S. Weiss, R. Voll, and L. Nitschke. 2010. Siglec-G regulates B1 cell survival and selection. J. Immunol 185:3277–3284.

Kanters, E., M.J. Gijbels, d.M. van, I, M.N. Vergouwe, P. Heeringa, G. Kraal, M.H. Hofker, and M.P. de Winther. 2004. Hematopoietic NF-kappaB1 deficiency results in small atherosclerotic lesions with an inflammatory phenotype. Blood 103:934–940.

Kanters, E., M. Pasparakis, M.J. Gijbels, M.N. Vergouwe, I. Partouns-Hendriks, R.J. Fijneman, B.E. Clausen, I. Forster, M.M. Kockx, K. Rajewsky, G. Kraal, M.H. Hofker, and M.P. de Winther. 2003. Inhibition of NF-kappaB activation in macrophages increases atherosclerosis in LDL receptor-deficient mice. J. Clin. Invest 112:1176–1185.

Kojima, Y., J.P. Volkmer, K. McKenna, M. Civelek, A.J. Lusis, C.L. Miller, D. Direnzo, V. Nanda, J. Ye, A.J. Connolly, E.E. Schadt, T. Quertermous, P. Betancur, L. Maegdefessel, L.P. Matic, U. Hedin, I.L. Weissman, and N.J. Leeper. 2016. CD47-blocking antibodies restore phagocytosis and prevent atherosclerosis. Nature 536:86–90.

Lekanne Deprez, R.H., A.C. Fijnvandraat, J.M. Ruijter, and A.F. Moorman. 2002. Sensitivity and accuracy of quantitative real-time polymerase chain reaction using SYBR green I depends on cDNA synthesis conditions. Anal. Biochem 307:63–69.

Libby, P., P.M. Ridker, and G.K. Hansson. 2011. Progress and challenges in translating the biology of atherosclerosis. Nature 473:317–325.

Maas, A., G.M. Dingjan, H.F. Savelkoul, C. Kinnon, F. Grosveld, and R.W. Hendriks. 1997. The X-linked immunodeficiency defect in the mouse is corrected by expression of human Bruton’s tyrosine kinase from a yeast artificial chromosome transgene. Eur J Immunol 27:2180–2187.

Matlung, H.L., K. Szilagyi, N.A. Barclay, and T.K. van den Berg. 2017. The CD47-SIRPalpha signaling axis as an innate immune checkpoint in cancer. Immunol Rev 276:145–164.

Miller, Y.I., and S. Tsimikas. 2013. Oxidation-specific epitopes as targets for biotheranostic applications in humans: biomarkers, molecular imaging and therapeutics. Curr. Opin. Lipidol 24:426–437.

Muggen, A.F., S.Y. Pillai, L.P. Kil, M.C. van Zelm, J.J. van Dongen, R.W. Hendriks, and A.W. Langerak. 2015. Basal Ca(2+) signaling is particularly increased in mutated chronic lymphocytic leukemia. Leukemia 29:321–328.

Myers, L.M., M.C. Tal, L.B. Torrez Dulgeroff, A.B. Carmody, R.J. Messer, G. Gulati, Y.Y. Yiu, M.M. Staron, C.L. Angel, R. Sinha, M. Markovic, E.A. Pham, B. Fram, A. Ahmed, A.M. Newman, J.S. Glenn, M.M. Davis, S.M. Kaech, I.L. Weissman, and K.J. Hasenkrug. 2019. A functional subset of CD8(+) T cells during chronic exhaustion is defined by SIRPalpha expression. Nat Commun 10:794.

Nitschke, L., R. Carsetti, B. Ocker, G. Kohler, and M.C. Lamers. 1997. CD22 is a negative regulator of B-cell receptor signalling. Curr. Biol 7:133–143.

Que, X., M.Y. Hung, C. Yeang, A. Gonen, T.A. Prohaska, X. Sun, C. Diehl, A. Maatta, D.E. Gaddis, K. Bowden, J. Pattison, J.G. MacDonald, S. Yla-Herttuala, P.L. Mellon, C.C. Hedrick, K. Ley, Y.I. Miller, C.K. Glass, K.L. Peterson, C.J. Binder, S. Tsimikas, and J.L. Witztum. 2018. Oxidized phospholipids are proinflammatory and proatherogenic in hypercholesterolaemic mice. Nature 558:301–306.

Ravandi, A., S.M. Boekholdt, Z. Mallat, P.J. Talmud, J.J. Kastelein, N.J. Wareham, E.R. Miller, J. Benessiano, A. Tedgui, J.L. Witztum, K.T. Khaw, and S. Tsimikas. 2011. Relationship of IgG and IgM autoantibodies and immune complexes to oxidized LDL with markers of oxidation and inflammation and cardiovascular events: results from the EPIC-Norfolk Study. J Lipid Res 52:1829–1836.

Sato-Hashimoto, M., Y. Saito, H. Ohnishi, H. Iwamura, Y. Kanazawa, T. Kaneko, S. Kusakari, T. Kotani, M. Mori, Y. Murata, H. Okazawa, C.F. Ware, P.A. Oldenborg, Y. Nojima, and T. Matozaki. 2011. Signal regulatory protein alpha regulates the homeostasis of T lymphocytes in the spleen. J. Immunol 187:291–297.

Smith, F.L., and N. Baumgarth. 2019. B-1 cell responses to infections. Curr Opin Immunol 57:23–31.

Smith, K.G., and M.R. Clatworthy. 2010. FcgammaRIIB in autoimmunity and infection: evolutionary and therapeutic implications. Nat Rev Immunol 10:328–343.

van den Berg, T.K., and T. Valerius. 2019. Myeloid immune-checkpoint inhibition enters the clinical stage. Nat Rev Clin Oncol 16:275–276.

Waffarn, E.E., C.J. Hastey, N. Dixit, Y. Soo Choi, S. Cherry, U. Kalinke, S.I. Simon, and N. Baumgarth. 2015. Infection-induced type I interferons activate CD11b on B-1 cells for subsequent lymph node accumulation. Nat Commun 6:8991.

Yamao, T., T. Noguchi, O. Takeuchi, U. Nishiyama, H. Morita, T. Hagiwara, H. Akahori, T. Kato, K. Inagaki, H. Okazawa, Y. Hayashi, T. Matozaki, K. Takeda, S. Akira, and M. Kasuga. 2002. Negative regulation of platelet clearance and of the macrophage phagocytic response by the transmembrane glycoprotein SHPS-1. J. Biol. Chem 277:39833–39839.

Yang, Y., J.W. Tung, E.E. Ghosn, L.A. Herzenberg, and L.A. Herzenberg. 2007. Division and differentiation of natural antibody-producing cells in mouse spleen. Proc. Natl. Acad. Sci. U. S. A 104:4542–4546.

